# Funders’ expectations for open science in cardiovascular research: A Scoping review of the largest cardiovascular research funders

**DOI:** 10.64898/2025.12.11.693828

**Authors:** Anna Catharina Vieira Armond, Al Mamoune Alaoui, David Moher, Jean Rouleau, Kelly D. Cobey

**Affiliations:** University of Ottawa Heart Institute, Ottawa, Canada; Centre for Journalology, Clinical Epidemiology Program, Ottawa Hospital Research Institute, Ottawa, Canada; School of Epidemiology and Public Health, Faculty of Medicine, University of Ottawa, Ottawa, Canada; Montreal Heart Institute, University of Montreal, Montreal, Canada

## Abstract

Open science practices, including data sharing, open access, and prospective study registration, have been increasingly recognized to improve transparency, reproducibility, and accessibility in research, yet their uptake and implementation by cardiovascular research funders is unclear. We conducted a scoping review of publicly available policies, guidance, and grant instructions from 12 members of the Global Cardiovascular Research Funders Forum to assess expectations, monitoring, and support for open science in cardiovascular research. We included 105 documents from 9 funders; no relevant documents were identified for 3 funders. Data sharing (75%) and open access (67%) were the most common mandates by funders, followed by prospective registration (50%). Requirements for other practices, including code sharing, use of reporting guidelines, preprints, and open peer review, were uncommon. Monitoring compliance was inconsistent, with many funders not specifying any mechanisms, even for widely required practices. Where available, support was most often provided through financial assistance, guidance, or infrastructure, particularly for open access, data sharing, and patient or public involvement. These findings suggest that while cardiovascular funders are engaging with open science, policies remain uneven in scope, monitoring, and support. Navigating the open science implementation gap in cardiovascular research will be essential to reap the benefits of transparency and innovation, only possible through the sharing of information and data.

## Introduction

Open science refers to a broad set of practices that aim to make scientific research more transparent, accessible, collaborative, and reproducible. While there is no single consensus on what open science entails, the practices include, but are not limited to, sharing research data, code, and materials; prospectively registering studies; publishing in open access; using reporting guidelines; and engaging in open peer review (1). Globally, open science has gained momentum as a strategy to foster research integrity, collaboration, and accelerate scientific discovery across disciplines, including biomedical research (1,2). Cardiovascular research, like many other fields, faces growing demands to improve the transparency and reproducibility of its research outputs (3). Prior audits show that most cardiology publications do not make data, protocols, or statistical analysis scripts publicly available (4), and present inconsistent reporting (5), echoing broader concerns about research waste and reproducibility (6,7). Open science practices offer a potential solution to these challenges, but their uptake remains variable (8), and implementation is often influenced by institutional and systemic factors, including the policies and expectations of research funders. These issues are especially pressing given the global burden of cardiovascular disease and the imperative to translate research findings efficiently into clinical care (9). Funders play a critical role in shaping research practices by setting requirements and providing incentives for grant recipients. By mandating or recommending open science practices, funders can promote a culture of transparency and reproducibility. Most previous analyses have focused on general biomedical funders or specific practices, such as data sharing (10), often neglecting the cardiovascular research landscape as a distinct domain with its own needs and challenges. For example, an international survey of 198 cardiovascular researchers (8) reported that many participants had limited formal training in open science: 52% learned largely on the job, and 36% had received no training at all. Respondents indicated that additional funding and institutional support were critical to adopting open science practices, with funders seen as key interest holders to influence their behaviours. Little is known about how cardiovascular research funders are addressing open science in their policies and guidance, and to what degree there is coordination internationally across cardiovascular funders.

To address this gap, we conducted a scoping review to identify and assess publicly available policies and guidance related to open science practices among funders that are members of the Global Cardiovascular Research Funders Forum (GCRFF), a global partnership of 12 major cardiovascular research funders (11). Our objectives were to (1) map the extent to which open science practices are addressed in these funders’ official documents, (2) classify their expectations (e.g., mandated, recommended), and (3) assess the types of support and monitoring mechanisms provided. By synthesizing this information, we aim to inform future efforts to align open science policies across funders and support their implementation in cardiovascular research.

## Methods

### Protocol and registration

We conducted a scoping review to identify and analyze publicly available policies and guidance related to open science practices from cardiovascular research funders. This review followed the JBI methodology for scoping reviews and is reported in accordance with the PRISMA Extension for Scoping Reviews (PRISMA-ScR) (12). The study protocol was registered prospectively (https://doi.org/10.17605/OSF.IO/9PZTA).

### Eligibility criteria

We included documents that addressed at least one open science practice (e.g., open access, open data, prospective registration, preprints) and were publicly available from cardiovascular research funders. Eligible document types included official policies, guidance documents, grant requirements, application instructions, and relevant web pages. Documents were included if written in English or if a reliable translation could be produced using DeepL Translate. Opinion pieces, blogs, and non-official funder communications were excluded.

### Information sources

We focused on members of the Global Cardiovascular Research Funders Forum (GCRFF), an international coalition of 12 major cardiovascular research funders:

1. American Heart Association (USA)
2. British Heart Foundation (UK)
3. Danish Heart Foundation
4. Dutch Heart Foundation (Hartstichting)
5. German Centre for Cardiovascular Research (DZHK)
6. Heart and Stroke Foundation of Canada
7. Institute of Circulatory and Respiratory Health (CIHR, Canada)
8. Leducq Foundation
9. National Heart Foundation of Australia
10. National Heart Foundation of New Zealand
11. National Heart, Lung and Blood Institute (NIH-NHLBI, USA)
12. Swiss Heart Foundation

We defined open science as an umbrella term encompassing practices that promote transparency, reproducibility, accessibility, and collaboration in research. Documents were included if they referred to at least one of the following practices: open access, open data, data management, open code, open materials, prospective registration, transparent reporting, reproducibility practices, preprints, citizen science (including patient and public involvement), open peer review, or researcher identifiers (e.g., ORCID). Table 1 presents the descriptions of each of the investigated practices.

**Table 1.** Open science practices descriptions.

| Open science practice | Description |
| --- | --- |
| Open access | The practice of making scholarly work freely available online, allowing anyone to access, read, and use it for lawful purposes without financial, legal, or technical barriers |
| Data sharing | The practice of making the underlying datasets used to generate research findings accessible to other researchers, policymakers, and the public, either openly or under controlled conditions |
| Code sharing | Code sharing in science refers to the practice of making the scripts, software, or computational workflows used in a study openly available to others. Sharing code allows peers to reproduce analyses, validate findings, and adapt existing methods for new research questions |
| Prospective registration | The practice of publicly registering a study's protocol, including research questions, outcomes, design, and planned analyses, before data collection or examination |
| Preprint | Preprints refer to scientific manuscripts or research findings that are made publicly available in advance of formal publication, typically referred to as a form of Green Open Access |
| Public and Patient Involvement | Public and Patient Involvement refers to the active partnership of patients, caregivers, and members of the public in the research process. Rather than being the subjects of research, they contribute to shaping research priorities, study design, conduct, and dissemination |
| Data Management Plans | Research data management (RDM) involves the systematic organization, storage, documentation, and preservation of research data throughout its lifecycle. DMPs are formal documents outlining how data will be collected, managed, shared, and preserved |
| Open peer review | Open peer review is an umbrella term for peer review models that aim to increase transparency in the evaluation of scholarly work. It can include practices such as revealing reviewer identities, publishing review reports alongside articles, allowing public comments, or enabling authors and reviewers to interact directly |
| Use of ORCID | Using a unique digital identifier for researchers that links their professional activities, publications, and datasets. ORCID enhances transparency, ensures proper attribution, and facilitates tracking of contributions across projects and platforms |
| Rigor and Reproducibility | Ensuring that research is conducted with robust, transparent, and unbiased methods so that results can be independently verified. Rigor refers to the strict application of scientific principles and methods, while reproducibility means that findings can be consistently obtained when experiments or analyses are repeated under the same conditions |
| Material sharing | Material sharing refers to the practice of making physical research resources, such as biological samples, cell lines, reagents, instruments, or other study materials, available to other researchers |
| Use of reporting guidelines | Tools or instructions designed to help authors transparently report research using explicit methods. These can take the form of checklists, flowcharts, or structured text to ensure clarity, completeness, and consistency in reporting |

We restricted our analysis to cardiovascular research funders and included only documents available on official websites, without date restrictions.

### Search strategy

Two reviewers independently conducted searches between January and April 2025. Searches were performed on funders’ websites using internal search tools and supplemented with Google queries combining funder names with targeted keywords (e.g., “open science,” “open data,” “data sharing,” “preprints,” “preregistration,” “reporting guideline,” “ORCID,” “patient involvement,” “open access,” and “reproducibility”). The first 100 Google hits per funder were screened for relevance.

### Data extraction and analysis

We developed a customized data extraction form and piloted it on five documents to ensure clarity and consistency. One reviewer extracted all data, and a second reviewer conducted quality control on all records. Discrepancies were resolved by consensus or with input from a third reviewer when necessary. Data extraction was performed using Airtable, a cloud-based database platform.

We extracted the following information:

1. Funder name and country
2. Document type (e.g., policy, grant call, guidance)
3. Year of publication or last update
4. Cardiovascular research focus (if specified)
5. Open science practices mentioned
6. Whether practices were required, recommended, mentioned without further detail, or not mentioned
7. Monitoring or compliance mechanisms
8. Forms of support (e.g., training, financial resources, infrastructure)

Some funders developed multiple documents corresponding to different funding opportunities, which varied in their expectations for open science practices. For example, one funder’s clinical trial program explicitly required data sharing, while its early-career investigator program only recommended it. To ensure consistency, we used an inclusive coding approach: if any eligible document from a funder mandated a given practice, we classified that practice as “Required” for the funder overall, even if other documents only recommended it or did not mention it. This approach reflects the presence of at least one formal requirement within the funder’s policy landscape.

Quantitative data were summarized descriptively using counts and proportions. For qualitative data, we conducted a thematic analysis following the approach by Braun and Clarke (13). Two reviewers independently reviewed and coded the extracted policy text, refined the codes through discussion, and organized them into overarching themes. Any disagreements were resolved with input from an additional reviewer. Thematic synthesis was conducted using Microsoft Excel (RRID: SCR_016137).

## Results

We identified 145 documents from 11 (of the 12) cardiovascular research funders, across 9 countries, in the GCRFF, based on searches on the funders’ websites and Google. For 1 funder, no potentially relevant documents could be located. After duplicates were removed, a total of 142 documents were assessed for eligibility. Of these, 38 documents were excluded for not meeting the inclusion criteria (e.g., documents lacked relevance to open science expectations or did not pertain to research funding). This resulted in 105 documents from 9 funders included in this review (Figure 1). The three funders where we did not locate relevant documents were: the Leducq Foundation, the National Heart Foundation of New Zealand, and the Swiss Heart Foundation.

**Figure 1.**
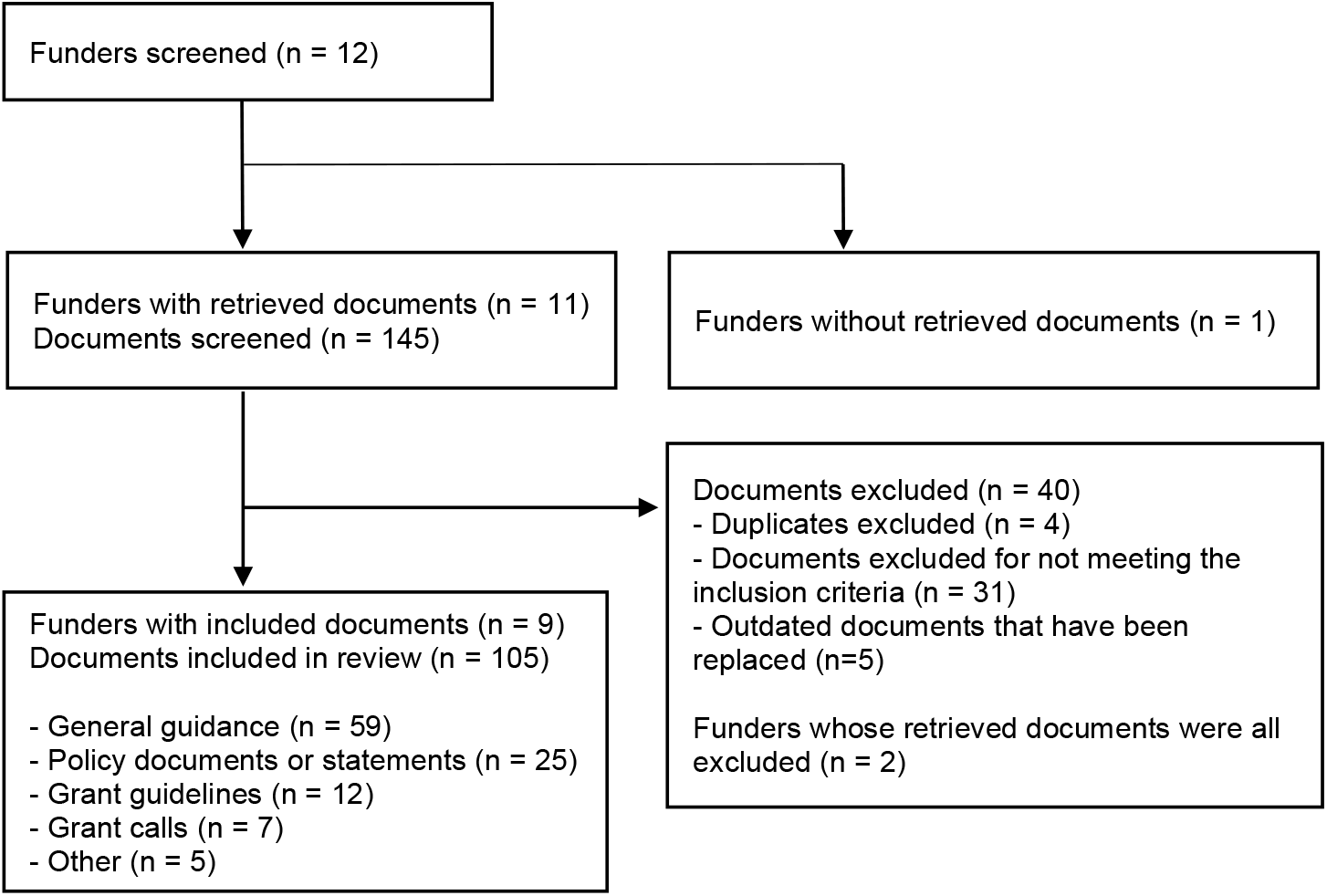
Flow-chart of funders and documents included in the scoping review.

Most documents were classified as General Guidance or Instructions (59, 55%), followed by Policy or Formal Statement (25, 23%), Grant Guidelines (12, 11%), Grant Calls (7, 6%), and Other (5, 5%). Where available, most documents were published or updated in 2024.

### Open science practices

We investigated 12 cardiovascular research funders for their expectations on open science practices. To calculate the proportion of funders addressing each open science practice, all 12 funders were included in the denominator. However, the three funders without any documents were not assigned classifications (e.g., “Required,” “Recommended,” or “Not Mentioned”). The extent to which various practices were required, recommended, mentioned, not mentioned, monitored, or supported is presented in Figure 1.

#### Open science expectations

The most commonly required practices were Data Sharing (9 funders, 75%) and Open Access (8 funders, 67%), followed by Prospective Registration (6 funders, 50%). Data Management Plans were required by 4 funders (33%) and recommended by 1 (8%). Similarly, Public and Patient Involvement was required by 4 funders (33%) and recommended by 1 (8%). Use of ORCID identifiers was required by 3 funders (25%), and Code Sharing was required by 2 funders (17%) and recommended by 1 (8%). Use of Reporting Guidelines was required by 2 funders (17%) (Figure 2).

**Figure 2.**
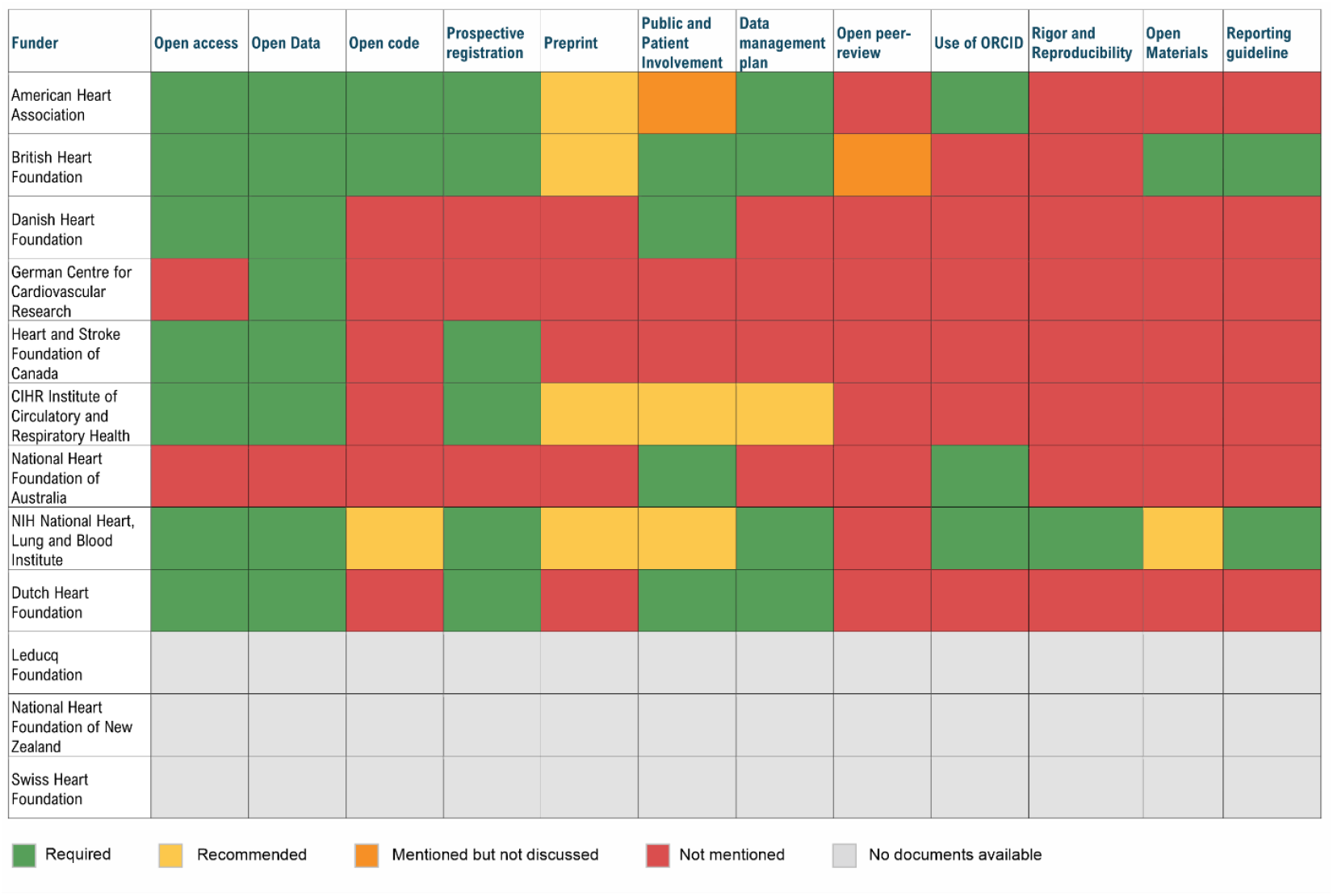
Funders’ expectations for open science practices.

Preprints were not required by any funders but were recommended by 4 (33%). Open Peer Review was rarely addressed, mentioned by only 1 funder (8%). Rigor and Reproducibility were explicitly required by just 1 funder (8%).

#### Monitoring and support of open science practices

The extent to which funders monitored compliance with open science practices and provided support varied across practice types (Tables 2 and 3). Among funders that required or recommended a given open science practice, monitoring mechanisms were inconsistently reported (Figure 2). For open access, 2 of 7 funders (29%) indicated that compliance was monitored, 3 (43%) provided unclear information, and 2 (29%) did not mention monitoring. For data sharing, 2 of 8 funders (25%) reported monitoring, 1 (13%) mentioned monitoring without further detail, and 5 (63%) did not mention monitoring. Monitoring was more frequent for data management plans (3 of 5, 60%), ORCID use (2 of 3, 67%), and rigor and reproducibility requirements (1 of 1, 100%). Prospective registration and public and patient involvement were monitored by one-third of relevant funders (2 of 6 each), whereas monitoring of code sharing was reported by 1 of 3 funders (33%). No monitoring mechanisms were identified for preprints, open peer review, material sharing, or use of reporting guidelines.

**Table 2.**
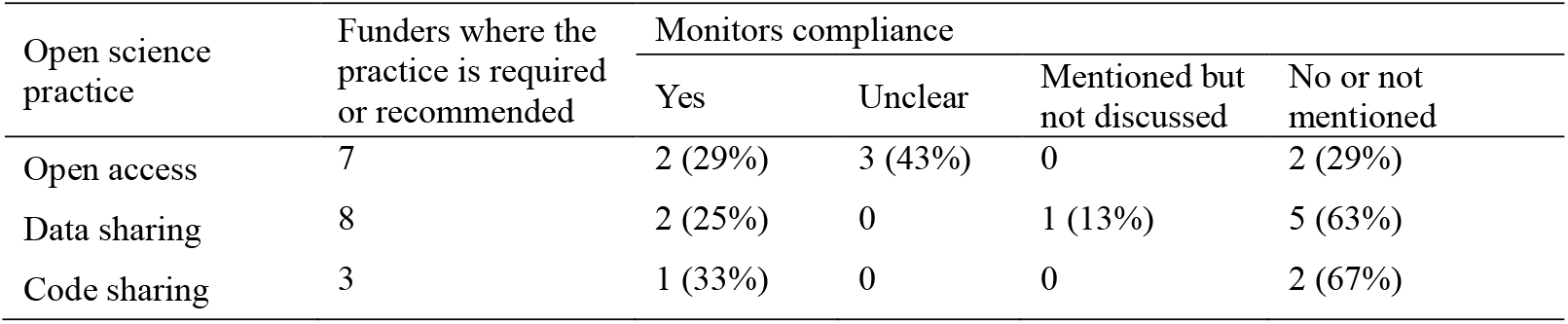
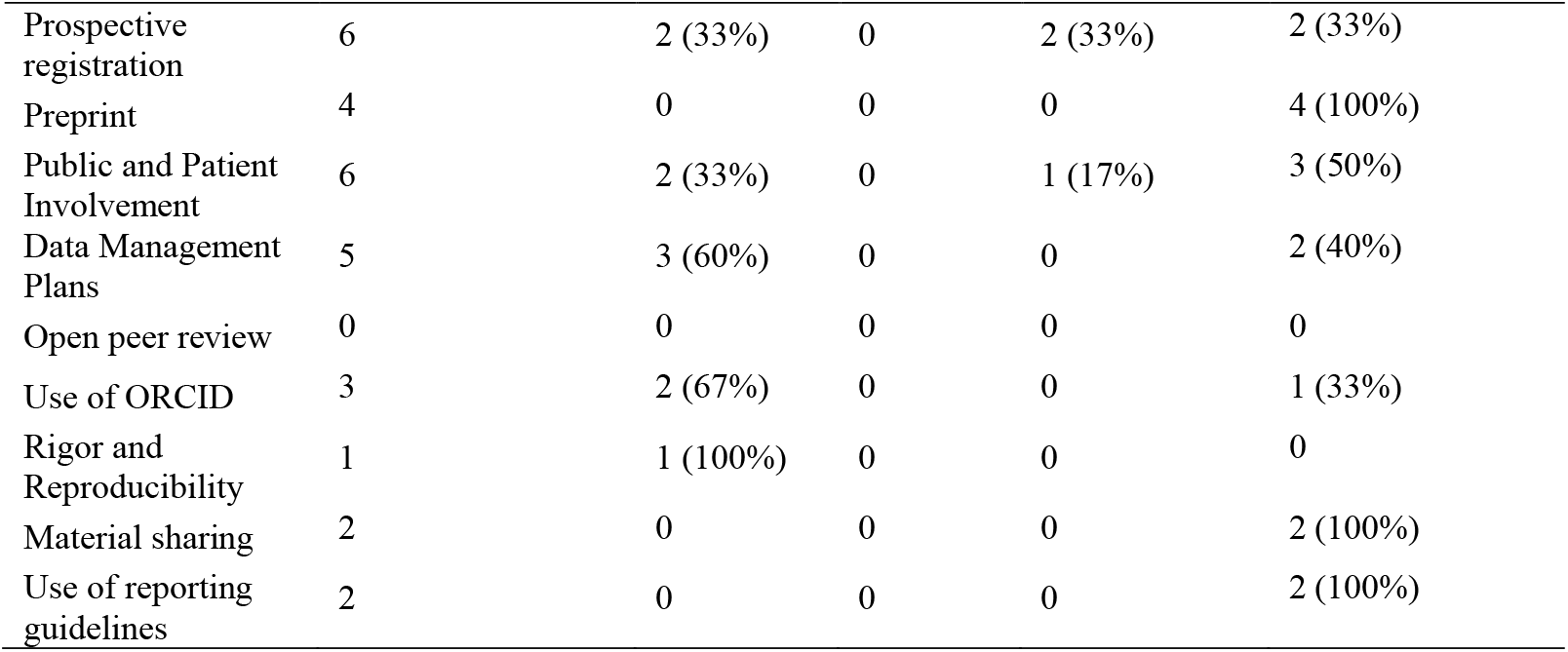
Monitoring of open science practices by cardiovascular research funders.

| Open science practice | Funders where the practice is required or recommended | Monitors compliance |  |  |  |
| --- | --- | --- | --- | --- | --- |
|  |  | Yes | Unclear | Mentioned but not discussed | No or not mentioned |
| Open access | 7 | 2 (29%) | 3 (43%) | 0 | 2 (29%) |
| Data sharing | 8 | 2 (25%) | 0 | 1 (13%) | 5 (63%) |
| Code sharing | 3 | 1 (33%) | 0 | 0 | 2 (67%) |
| Prospective registration | 6 | 2 (33%) | 0 | 2 (33%) | 2 (33%) |
| Preprint | 4 | 0 | 0 | 0 | 4 (100%) |
| Public and Patient Involvement | 6 | 2 (33%) | 0 | 1 (17%) | 3 (50%) |
| Data Management Plans | 5 | 3 (60%) | 0 | 0 | 2 (40%) |
| Open peer review | 0 | 0 | 0 | 0 | 0 |
| Use of ORCID | 3 | 2 (67%) | 0 | 0 | 1 (33%) |
| Rigor and Reproducibility | 1 | 1 (100%) | 0 | 0 | 0 |
| Material sharing | 2 | 0 | 0 | 0 | 2 (100%) |
| Use of reporting guidelines | 2 | 0 | 0 | 0 | 2 (100%) |

**Table 3.** Support provided by cardiovascular research funders for each open science practice.

| Open science practice | Funders where the practice is required or recommended | Provides support | Provided support |  |  |  |  |
| --- | --- | --- | --- | --- | --- | --- | --- |
|  |  |  | Cover related costs | Provides instructions or guiding materials | Provides infrastructure | Provides formal or informal training | Not mentioned |
| Open access | 7 | 6 (86%) | 6 (86%) | 3 (43%) | 2 (29%) | 1 (14%) | 3 (43%) |
| Data sharing | 8 | 4 (50%) | 4 (50%) | 3 (38%) | 3 (38%) | 1 (13%) | 4 (50%) |
| Code sharing | 3 | 2 (67%) | 0 (0%) | 1 (33%) | 2 (67%) | 0 (0%) | 1 (33%) |
| Prospective registration | 6 | 4 (67%) | 1 (17%) | 4 (67%) | 0 (0%) | 1 (17%) | 2 (33%) |
| Preprint | 4 | 3 (75%) | 0 (0%) | 3 (75%) | 0 (0%) | 0 (0%) | 1 (25%) |
| Public and Patient Involvement | 6 | 4 (67%) | 3 (50%) | 4 (67%) | 0 (0%) | 1 (17%) | 2 (33%) |
| Data Management Plans | 5 | 4 (80%) | 1 (20%) | 4 (80%) | 0 (0%) | 2 (40%) | 1 (20%) |
| Open peer review | 0 | - | - | - | - | - | - |
| Use of ORCID | 3 | 1 (33%) | 0 (0%) | 1 (33%) | 0 (0%) | 0 (0%) | 2 (67%) |
| Rigor and Reproducibility | 1 | 1 (100%) | 0 (0%) | 1 (100%) | 0 (0%) | 1 (100%) | 0 (0%) |
| Material sharing | 2 | 0 (0%) | 0 (0%) | 0 (0%) | 0 (0%) | 0 (0%) | 2 (100%) |
| Use of reporting guidelines | 2 | 1 (50%) | 0 (0%) | 1 (50%) | 0 (0%) | 0 (0%) | 1 (50%) |

Support mechanisms varied widely across practices (Table 3). Financial assistance to cover related costs, through dedicated funding or fee reimbursements, was most frequently offered for open access (86%), data sharing (50%), and public and patient involvement (50%), reflecting the resource demands of these practices. Guiding materials were widely provided across several practices, including open access (43%), data sharing (38%), prospective registration (67%), and data management plans (80%). Infrastructure, such as data repositories, was provided for data and code sharing (38% and 67%, respectively), and open access (29%). Formal or informal training was less commonly reported overall, but was most commonly provided for data management plans (40%), prospective registration, and public and patient involvement (17%).

## Discussion

This scoping review provides a comprehensive assessment of open science expectations among 12 major cardiovascular research funders globally. Our findings show that while many funders have incorporated some key open science practices, formal requirements remain unevenly applied across the range of open science activities. Notably, data sharing and open access were the most frequently required practices, reflecting broad recognition of their central role in fostering accessibility and transparency of science. However, practices such as code sharing, use of reporting guidelines, open peer review, and preprints received far less attention, suggesting important gaps in funder policies.

Monitoring of compliance with open science policies was inconsistent and often lacking, even for widely required practices. Without (continuous) monitoring, it is likely difficult for funders to use data to better disseminate and implement open science mandates and recommendations among their grantees. For example, fewer than one-third of funders with requirements for open access publishing or data sharing actively monitored adherence. Monitoring was somewhat more common for data management plans (60%) and the use of ORCID (67%), which may reflect their stronger ties to administrative infrastructure or grant reporting mechanisms. The lack of monitoring mechanisms for practices such as preprints, open peer review, and material sharing indicates a potential disconnect between funder expectations and enforcement, raising questions about the effectiveness of current policies in promoting sustained behavior change among researchers.

Despite variable monitoring, many funders provide important support mechanisms to facilitate implementation. Financial assistance to cover related costs was more frequent for open access (86%), data sharing (50%), and public and patient involvement (50%), highlighting funders’ recognition of cost barriers for these practices. Guiding materials and infrastructure, including repositories and technical platforms, were also frequently made available for data sharing, code sharing, and data management plans, supporting researchers in meeting policy requirements.

However, formal or informal training opportunities were less consistently reported, highlighting a potential area for growth. Given that prior surveys have identified a lack of training and resources as key barriers to open science (8) uptake in cardiovascular research, increasing capacity-building initiatives may be critical to achieving more widespread adoption.

This limited and uneven adoption of open science reflects a lack of coordination that could limit the global cardiovascular research community from fully benefiting from open science. Given that cardiovascular research is inherently international, investigators in any single region rely on access to publications, data, and protocols produced worldwide. Without harmonized policies and consistent enforcement, the potential for open science to accelerate discovery and improve patient outcomes is undermined.

## Limitations

This study has several strengths, including its novel focus on cardiovascular research funders, a systematic and rigorous approach to identifying and analyzing policies, and an assessment of a broad range of open science practices within a single review. However, some limitations must be acknowledged. Our search was limited to publicly available documents, excluding those only accessible behind password-protected platforms such as grant application portals. Many funders make their requirements available only at these later stages. We did not contact funders to verify the completeness of the documents, including cases where no documents were identified. This is a limitation of our study. Nonetheless, we consider it essential that open science expectations be made publicly available and easily accessible. Clear communication of these requirements before application is crucial to ensure transparency and to allow researchers to prepare adequately.

Additionally, our searches were conducted using keywords in both English and French, and we included documents for which reliable translations could be obtained using automatic tools such as browser-based translators available on funders’ websites. Despite these efforts, documents published exclusively in other languages or lacking accessible translations may have been missed and underrepresented in our review.

## Conclusion

In conclusion, our findings suggest that cardiovascular funders are engaging with open science but face challenges in translating policies into practice. The observed gaps in monitoring and support highlight opportunities for funders to strengthen their policies by integrating clearer compliance mechanisms and expanding resources for researchers. Coordinated efforts among funders, aligned with evolving best practices and researcher needs, will be essential to fostering a culture of transparency and reproducibility that can accelerate discovery and ultimately improve cardiovascular research findings.

Future work should explore how funder policies interact with institutional and researcher-level factors to influence open science behaviors in cardiovascular research. For example, if a funder mandated data sharing but the grantee’s institution did not, this might be seen as a barrier to implementing the practice. It is likely that better communication between funders and research-performing organizations will foster greater uptake and implementation of open science practices. Additionally, qualitative research engaging key interest holders could help identify practical barriers and facilitators to policy implementation, informing more tailored and effective interventions. As open science continues to evolve, funders’ roles as leaders and enablers will remain critical in shaping a more open, collaborative, and trustworthy cardiovascular research ecosystem. If cardiovascular research wishes to reap the benefits afforded by open science, funders will want to ensure that they set policies, resources, and provide guidance for researchers to comply, and monitor success.

## Author Contributions

**Conceptualization:** All authors; **Methodology:** All authors; **Formal Analysis:** ACVA, AMA, KDC; **Investigation:** ACVA, AMA; **Data Curation:** ACVA, AMA; **Writing-Original Draft:** ACVA; **Writing-Review & Editing:** All authors**; Supervision:** KDC; **Project Administration:** ACVA, KDC; **Funding Acquisition:** KDC, DM, JR.

## Data availability statement

All data and materials underlying this study are publicly available on the Open Science Framework (OSF) at: https://doi.org/10.17605/OSF.IO/9JEFW.

